# Non-canonical RNA substrates of Drosha lack many of the conserved features found in primary microRNA stem-loops

**DOI:** 10.1101/2023.03.30.535007

**Authors:** Karen Gu, Lawrence Mok, Matthew J. Wakefield, Mark M.W. Chong

## Abstract

The RNase III enzyme Drosha has a central role in microRNA (miRNA) biogenesis, where it is required to release the stem-loop intermediate from primary (pri)-miRNA transcripts. However, it can also cleave stem-loops embedded within messenger (m)RNAs. This destabilizes the mRNA causing target gene repression and appears to occur primarily in stem cells. While pri-miRNA stem-loops have been extensively studied, such non-canonical substrates of Drosha have yet to characterized in detail. In this study, we employed high-throughput sequencing to capture all polyA-tailed RNAs that are cleaved by Drosha in mouse embryonic stem cells (ESCs) and compared the features of non-canonical versus miRNA stem-loop substrates. First, mRNA substrates are less efficiently processed than miRNA stem-loops. Sequence and structural analyses revealed that these mRNA substrates are also less stable and more likely to fold into alternative structures than miRNA stem-loops. Moreover, they lack the sequence and structural motifs found in miRNAs stem-loops that are required for precise cleavage. Notably, we discovered a non-canonical Drosha substrate that is cleaved in an inverse manner, which is a process that is normally inhibited by features in miRNA stem-loops. Our study thus provides valuable insights into the recognition of non-canonical targets by Drosha.

## INTRODUCTION

MicroRNAs (miRNAs) are small RNAs, ∼22 nucleotides (nt) in length, that play a crucial role in regulating gene expression (1,2). Mature miRNAs are generated from stem-loop structures embedded within primary (pri)-miRNA transcripts (3,4). The biogenesis of most miRNAs requires two RNase III enzymes, Drosha and Dicer, that process precursors in a stepwise manner (3,4). Drosha interacts with a dimer of the RNA-binding protein Dgcr8 to form the microprocessor complex (5-7). In the nucleus, the microprocessor binds to and cleaves the stem-loop structure in the pri-miRNA, releasing a precursor (pre)-miRNA stem-loop intermediate from the flanking single-stranded RNA (ssRNA) segments (3,4). The pre-miRNA is exported to the cytoplasm where it is further processed by Dicer. Dicer cleaves 21–22 nt from the Drosha cleavage sites, generating a miRNA duplex (3,4). This miRNA duplex is then bound by an Argonaute protein and one (passenger) strand quickly dissociates while the other (guide) strand remains with the Argonaute to forms the core of the miRNA-induced silence complex (miRISC) (8). The mature miRNA guides the complex to target mRNAs that have complementarity to its seed region (2–7 nt from the 5’ end of the miRNA) to induces gene repression (2,3).

Accurate processing of miRNA precursors is crucial because even a single nucleotide shift in the seed sequence can result in miRISC binding to an entirely different set of mRNA targets. Because cleavage of the pre-miRNA by Dicer is entirely based on the position of Drosha cleavage (9), precise cleavage by Drosha is particularly critical. Pri-miRNAs have been shown to contain multiple sequence and structure motifs that ensure accurate Drosha cleavage.

Structurally, a miRNA stem-loop has an extensively base-paired stem of ∼35±1 bp (10). Two strands of the stem are connected by a ssRNA apical loop and flanked by unstructured ssRNA segments at the base. The microprocessor binds to the dsRNA stem, with Dgcr8 positioned towards the apical loop and Drosha towards the base with the catalytic site located ∼11 bp away from the basal junction (11,12). Multiple motifs have been identified on miRNA stem-loops that are crucial for ensuring accurate cleavage. These motifs include the basal UG motif, the apical UGU/GUG (UGU) motif, the CNNC motif, the mGHG motif, and the midMW motif (10,13,14). These motifs appear to be redundant as the presence of a single motif is sufficient to enable efficient and accurate cleavage by Drosha.

In addition to pri-miRNAs, Drosha can also cleave mRNAs that harbor stem-loops resembling those in pri-miRNAs (15). However, the stem-loop products from this process rarely enter the miRNA biogenesis pathway. Instead, this cleavage primarily destabilizes the mRNA, leading to the repression of target gene expression. Direct Drosha cleavage-mediated gene repression has been demonstrated for the *Dgcr8* (16-18), *Myl9, Todr1* (19), *Ngn2* (20), and *Nfib* (21) mRNAs. Drosha cleavage of the *Dgcr8* mRNA has been observed in many cell types. Because Dgcr8 is part of the microprocessor, this cleavage is thought to function as an autoregulatory mechanism. Drosha cleavage of the other mRNAs has only been observed in pluripotent stem cells and cleavage plays a crucial role in maintaining the pluripotency of these cells (15).

High-throughput sequencing suggests that there may be dozens of mRNAs that are cleaved by Drosha (22,23). While the features of pri-miRNA stem-loops have been extensively studied, the Drosha-targeted mRNAs have not been as well characterized. In this study, we employed high-throughput sequencing to capture Drosha-cleaved polyA RNA in mouse embryonic stem cells (ESCs) and characterized the features of these non-canonical stem-loop substrates of this enzyme. We show that while Drosha cleaved pri-miRNA stem-loops and mRNA stem-loops may appear similar at a superficial level, there are fundamental differences between these two groups of substrates.

## MATERIAL AND METHODS

### Mouse ESCs culturing

Conditional LoxP-flanked *Drosha* ^*fl/fl*^ *Gt(ROSA)26Sor*^*CreERT2*^ ESCs have previously been described, which were originally derived from blastocysts obtained from time-mated *Drosha* ^*fl/fl*^ *Gt(ROSA)26Sor*^*CreERT2*^ mice (24).

Conditional LoxP-flanked *Dicer*^*fl/fl*^ ESCs were derived from blastocysts obtained from *Dicer1*^*fl/fl*^ mice (25). In brief, embryonic day 3.5 blastocysts were flushed from the uterine horns of time-mated female mice. Individual blastocysts were then deposited into wells containing Mitomycin C-inactivated mouse embryonic fibroblasts as feeder cells and cultured in KnockOut DMEM (Gibco) supplemented with 20% Knockout Serum Replacement (Gibco), non-essential amino acids (Gibco), 0.1 mM β2-mercaptoethanol and 10^3^ U/mL LIF (Peprotech). The wells were monitored daily for outgrowth from the blastocysts. Upon outgrowth, the well was trypsinized into single cell suspension and transferred into a larger well for picking of individual colonies.

All ESCs were cultured in KnockOut DMEM, supplemented with 10% heat-inactivated fetal bovine serum (GE Healthcare Life Sciences), 5% KnockOut Serum Replacement (Gibco), 1% sodium pyruvate (Gibco), 1% non-essential amino acids, 1% penicillin-streptomycin-glutamine (Gibco), 0.1 mM 2-mercaptoethanol and 10^3^ U/mL LIF on a Mitomycin C-inactivated mouse embryonic fibroblasts.

Deletion of the floxed Drosha allele was achieved by adding 100 nM 4-hydroxytamoxifen (Sigma-Aldrich) to the *Drosha* ^*fl/fl*^ *Gt(ROSA)26Sor*^*CreERT2*^ ESCs for 72h. The medium was then replaced (without 4-hydroxytamoxifen) for a further 48h to allow for depletion of Drosha-dependent miRNAs before analysis.

Deletion the floxed Dicer allele was achieved by transducing the *Dicer*^*fl/fl*^ ESCs with a Cre-expressing retrovirus (26). The virus also contained a GFP reporter that allowed for sorting of transduced cells. GFP^+^ ESCs were sorted 3 days after transduction, then cultured for a further 2 days before analysis.

### Quantitative (q)RT-PCR

Total RNA was extracted from the cells using TRIsure (Bioline) following the manufacturer’s instruction. For measuring mRNA expression, 1 μg total RNA was reverse transcribed with 50 ng random hexamers using M-MuLV reverse transcriptase (NEB). 1/20^th^ of the resulting cDNA was used then analyzed by qRT-PCR using GoTaq qPCR master mix (Promega). The following primer pairs were used:

5’-GACGACGACAGCACCTGTT-3’ and 5’-GATAAATGCTGTGGCGG-ATT-3’ for Drosha; 5’-TCTGCAGGCTTTTACACACG-3’ and 5’-CAGCCAATGATGCAAA-GATG-3’ for Dicer; 5’-CACAGCTTCTTTGCAGCTCCTT-3’ and 5’-CGTCATCCATGGCGAAC-TG-3’ for β-actin.

Taqman miRNA assays (Thermo Fisher) were used to quantify the expression of mature miRNAs and U6 snRNA (control). For each target, 10 ng total RNA was reverse transcribed with its specific reverse transcription primer using 25 U Multiscribe reverse transcriptase. qRT-PCR then performed on 1/15^th^ of the cDNA with Taqman universal PCR master mix and 1X Taqman miRNA assay mix (miR-16 assay ID: 000391; miR-191 assay ID: 002299; miR-320 assay ID: 002277; miR-484 assay ID: 001821; U6 snRNA assay ID: 001973).

### Degradome sequencing library construction

Degradome sequencing (Degradome-seq) libraries were constructed based on a previously described protocol (22). In brief, polyA-tailed RNA was isolated from 75 μg of total RNA using the Dynabeads mRNA direct purification kit (Thermo Fisher) following the manufacturer’s instructions. An RNA linker (5’-CACGACGCUCUUCCGAUCU-3’) was ligated to the polyA RNA with T4 RNA ligase (Thermo Scientific) to capture RNAs with a 5’ monophosphate (5’P), a hallmark of RNase III cleavage. The ligation products were then purified with the Dynabeads mRNA direct purification kit and reverse transcription was performed using Superscript III reverse transcriptase (Invitrogen) and random hexamer primer attached to reverse adaptor sequence (5’-AGACGTGTGCTCTTCCGATCNNNNNN-3’). The cDNA was cleared of RNA by treating with RNase H (NEB). Half the cDNA was subjected to 2^nd^ strand synthesis using Phusion high-fidelity DNA polymerase (NEB) with primers to the 5’ RNA linker and reverse adaptor (5’-ACACTCTTTCCCTACACGACGCTCTTCCGATCT-3’ and 5’-GTGACTGGAGTTCAGACGTGTGCTCTTCCGATC-3’) using the cycling conditions: 98°C for 1 minute, (98°C for 30 seconds, 58°C for 30 seconds, 72°C for 1 minute) x7 cycles, and 72°C for 5 minutes. The resulting cDNA was resolved on a Low Melting Point agarose gel (Scientifix) and cDNA corresponding to 200–400 bp were gel purified. This cDNA library was then PCR barcoded with PCR Primer 1.0 (5’-AATGATACGGCGACCACCGAGATCTACACTCTTTCCCTACACGACG-3’) and a sample-specific barcode primer (5’-CAAGCAGAAGACGGCATACGAGATNNNNNNGTGACTGGAGTTCAGACGTG-3’) using MyTaq Red Mix (Bioline) for 15 cycles. The resulting cDNA libraries were sequenced on the NextSeq 500 platform (Illumina) for 75 cycles in single-end high-output mode at the Australian Genome Research Facility.

### Sequence processing and alignment of Degradome-seq libraries

Processing and alignment of Degradome-seq libraries were performed on the Galaxy Australia platform (27). The raw reads were processed using the Trimmomatic program (28) (version 0.38.0) to remove Illumina platform-specific adaptor sequences and low-quality bases (below 20 across 4 nt). The processed reads were then aligned to the mouse reference genome GRCm38 (mm10) using RNA STAR with default parameters (29) (version 2.7.8a).

### Degradome-seq analysis pipeline

The depth of Degradome-seq reads at each locus was determined by counting the number of reads starting at that locus. The raw counts were normalized as the reads per million (RPM). The depth of read at the 5’ end was denoted by *C5*. For each genomic locus i, we generated a counting vector *C*_*i*_ by

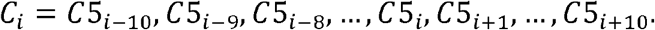

The counting vectors at known Drosha cleavage sites in annotated pri-miRNAs were used as positive training dataset. A set of 1000 loci was randomly selected from the rest of the genome as a negative training dataset. These positive and negative training datasets were used to train a generalized linear model using the *cv*.*glmnet* function in the R package *glmnet* (30), with the following parameters: family = “binomial”, type.measure = “class”, nfolds = 10. The training error was accessed by sensitivity, specificity, and the area under the curve (AUC). The AUC was calculated using the R package *pROC* (31) (version 1.18.0). The sensitivity and specificity were calculated using the formulas below:

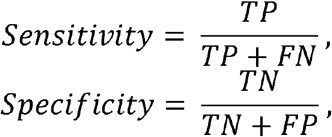

where *TP* is true positive, *FN* is false negative, *TN* is true negative, and *FP* is false positive. We considered the model trained successfully if it had sensitivity > 0.9, specificity > 0.9, and AUC > 0.9. We then applied the model to the rest of the genomic loci to identify those with a similar stacking pattern (i.e., pile-up site) to the known pri-miRNAs. The step between randomly selecting 1000 negative samples and applying the model to the rest of the genomic loci was repeated until the datasets were classified 1000 times. A locus was considered a true pile-up site if it was classified as a pile-up site in more than 900 iterations in both control ESC replicates.

To determine whether a pile-up site was dependent on Drosha or Dicer, we performed differential pile-up analysis between Drosha deficient and control ESCs or Dicer deficient and control ESCs using the *edgeR* package (32) (version 3.26.1). Raw read counts starting at each genomic locus were normalized by the trimmed mean of M values (TMM), and the dispersion was estimated by calling the function *estimateDisp()*. The testing of differential pile-ups was performed by calling functions *glmQLFit()* and *glmQLFTest()*. A genomic locus with a log_2_ fold change (log_2_FC) < −1.5 and false discovery rate (FDR) < 0.05 were considered significant.

### Sequence processing and alignment of small RNA sequencing libraries

Small RNA sequencing (sRNA-seq) libraries used in this study are retrieved from the NCBI Gene Expression Omnibus (GEO) database (Supplementary Table S1) and the processing and alignment of reads were performed on the Galaxy Australia platform (27). The quality of the reads and the representation of the sequencing adapters were accessed by the FastQC program (version 0.72), and the reported adapter sequences were trimmed using cutadapt (33) (version 1.16). The processed reads were collapsed into unique reads using an in-house. The unique reads were then aligned to the mouse reference genome GRCm38 (mm10) using RNA STAR with default parameters (29) (version 2.7.8a).

### Identification of Drosha-cleaved stem-loops

Positions identified as Drosha-dependent cleavage sites in the Degradome-seq analysis were denoted as *D*_*i*_. Reads starting or ending between *D*_*i*−100_ and *D*_*i*−25_ or between *D*_*i+*25_ and *D*_*i+*100_were collected. If the sRNA-seq reads stack at one of the termini and the depth of that terminus is > 1 RPM (denoted as *S*_*j*_), the RNA sequence between *S*_*j*−15_ and *D*_*i+*15_ or between *D*_*i*−15_ and *S*_*j*+15_ was used to predict the secondary structure using *RNAfold* program in the *ViennaRNA* package (34) (version 2.4.17) with the following option: RNAfold -p -d2 --noLP --MEA. The 3’ overhang was calculated in the structure generated by *RNAfold*. The structure was considered an authentic Drosha-cleaved stem-loop if *S*_*j*_ and *D*_*i*_ form a 3’ 2±2 nt overhang.

### Analysis of stem-loop thermodynamic properties

The length of the stem-loop was determined as the distance between *D*_*i*_ and *S*_*j*_ with a 15 nt flanking sequence. The base pairing frequency and maximum stacking of base pairing were normalized to the length of the stem-loop. The base pair composition was normalized to the total number of base pairs in the upper- or lower-stem.

### Analysis of stem-loop structural diversity

The ensemble diversity of the stem-loop was obtained from the *RNAfold* output using the option described above. The positional entropy of the stem-loops was calculated using script *mountain*.*pl* in the *ViennaRNA* package (34) (version 2.4.17).

### Information bits plot of structural features

The secondary structure between *D*_*i*_ and *S*_*j*_ with a 25 nt flanking sequence was predicted using the *RNAfold* program with the following option: RNAfold -p -d2 --noLP -C. The position of paired bases in secondary structure between *D*_*i*_ and *S*_*j*_ with a 15 nt flanking sequence was used as a constraint to inform the prediction. This ensured that paired bases remained paired when incorporated with longer flanking segments.

An in-house script was used to determine whether a position forms a base pair, a bulge, an internal loop, a terminal loop, or flanking segments. The resulting data was used to generate information bits plots with the python package *Logomaker* (35) (version 0.8).

### Sequence Logo

The sequence logo was generated using the python package *Logomaker* (35) (version 0.8) and the sequence between *D*_*i*_ and *S*_*j*_ with a 25 nt flanking sequence.

### Statistical analysis

Two-tailed *t*-tests were used to perform statistical analysis on the qPT-PCR data, and two-tailed unequal variance *t*-tests were used to analyze the differences between miRNA and non-miRNA stem-loop features. Results were considered statistically significant at *p* < 0.05.

## RESULTS

### Capturing Drosha-cleaved polyA RNAs by Degradome-seq

Drosha is known to cleave mRNA in mouse ESCs (22). To characterize the non-canonical RNAs that are directly cleaved by Drosha in mouse ESCs, we employed Degradome-seq to capture polyA-tailed RNAs with a 5’P, a hallmark of Drosha-mediated cleavage (Figure 1A). The Degradome-seq libraries were constructed essentially as described in (22) (Supplementary Figure S1). PolyA RNA was first extracted, and an RNA linker was ligated to RNAs with a 5’P. The RNAs were then reverse transcribed with a random hexamer attached to second (DNA) reverse linker sequence. PCR with primers against the 5’ and reverse linkers amplified fragments that have been endonucleolytically cleaved. To determine which cleavage sites are dependent on Drosha, we compared the sites between Drosha-deficient and control ESCs. These cells harbor a LoxP-flanked *Drosha*^*fl/fl*^ allele and CreERT2 knocked into the *Rosa26* locus (24). Addition of 4-hydroxytamoxifen ablated Drosha expression and consequently the expression of canonical miRNAs, like miR-16 and miR-191, while the expression of Dicer was unaffected (Figure 1B and C). The expression of the non-canonical miRNAs that are independent of Drosha, like miR-320 and miR-484 (36), were unaffected (Figure 1C). Two independent libraries were generated from both Drosha deficient and control cells, each resulting in ∼20 million reads mapping uniquely to the mouse reference genome GRCm38 (mm10; Supplementary Table S1).

**Figure 1.**
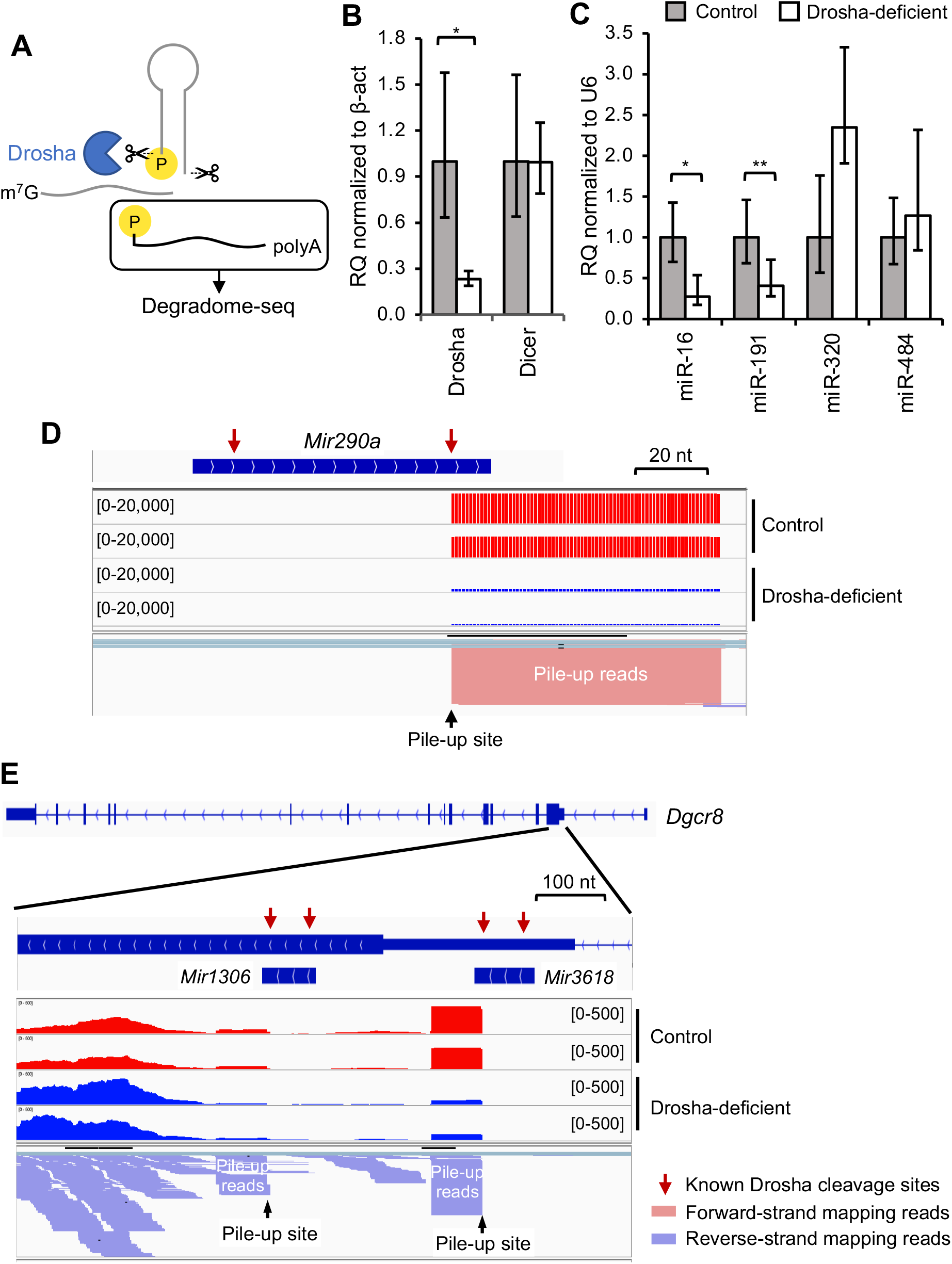
Capturing Drosha-cleaved RNA fragments by Degradome-seq. **(A)** Degradome-seq captures polyA-tailed RNA with a 5’P, which is the hallmark of Drosha cleavage. **(B)** qRT-PCR analysis for Drosha and Dicer mRNA levels in Drosha-deficient ESCs, normalized to β-actin. The data shows the mean ± S.E.M of 2 replicates. **(C)** Taqman qRT-PCR analysis for expression of select mature canonical miRNAs (miR-16 and miR-191) and non-canonical miRNAs (miR-320 and miR-484) in Drosha-deficient ESCs, normalized to U6 snRNA. The data shows the mean ± S.E.M of 2 replicates. **(D)** Example of Degradome-seq reads mapping to a miRNA gene (*Mir290a)* locus. **(E)** Degradome-seq reads mapping to the stem-loops within the 5’UTR and CDS of *Dgcr8*. Both these stem-loops are also annotated as miRNAs. In **(D)** and **(E)**, gene/miRNA genomic positions are indicated in the top panel, with identified cleavage sites indicted by the read arrows. Read depth is shown in next four tracks, with the data range for each track indicated in square brackets. The last track shows a collapsed view of aligned reads. Statistical testing in **(B)** and **(C)**: * *p* < 0.05; ** *p* < 0.01 (*t*-test).

We first validated the Degradome-seq libraries by examining reads that map to annotated canonical miRNA loci. As expected, these reads exhibited a homogenous 5’ stack precisely at the Drosha cleavage site on the 3’ arm of the stem-loop (Figure 1D). This detection of the 3’ cleavage site is expected because protocol captures polyA-tailed RNA fragments (Figure 1A). These stacked reads are referred to as “pile-up reads”, with the homogenous 5’ end referred to as the “pile-up site”. The depth of the pile-up site at the example *Mir290a* was reduced in Drosha-deficient ESCs, confirming that cleavage is Drosha-mediated (Figure 1D).

An analogous stacking pattern was observed at other mRNA targets of Drosha. For example, Drosha-dependent pile-up sites are observed at the two annotated stem-loops in *Dgcr8*, one in the 5’ untranslated region (UTR) and one in the coding domain sequence (CDS; Figure 1E). These results demonstrate the effectiveness of our Degradome-seq libraries for capturing Drosha cleavage sites in both pri-miRNAs and mRNA substrates.

Taking advantage of the clear stacking pattern in known substrates of Drosha, we developed a bioinformatics pipeline to systematically identify all Drosha-dependent cleavage events (Figure 2A). We first counted the depth at the start of reads for each locus. These counts were used to train a classification algorithm to learn the typical stacking pattern at known Drosha cleavage sites in pri-miRNAs. The successfully trained algorithm was then applied to the remaining loci to assess whether they exhibit a similar stacking pattern. This method identified 3632 pile-up sites in control cells, with 913 of these significantly decreased in Drosha-deficient cells (including pri-miRNAs; Figure 2B; Supplementary Table S2). The pipeline successfully captured the cleavage of canonical pri-miRNAs, such as the *Mir290∼295* family, while excluding Drosha-independent pri-miRNAs, such as *Mir5099*. These findings indicate that our pipeline successfully identifies Drosha-dependent cleavage of polyA RNAs in ESCs.

**Figure 2.**
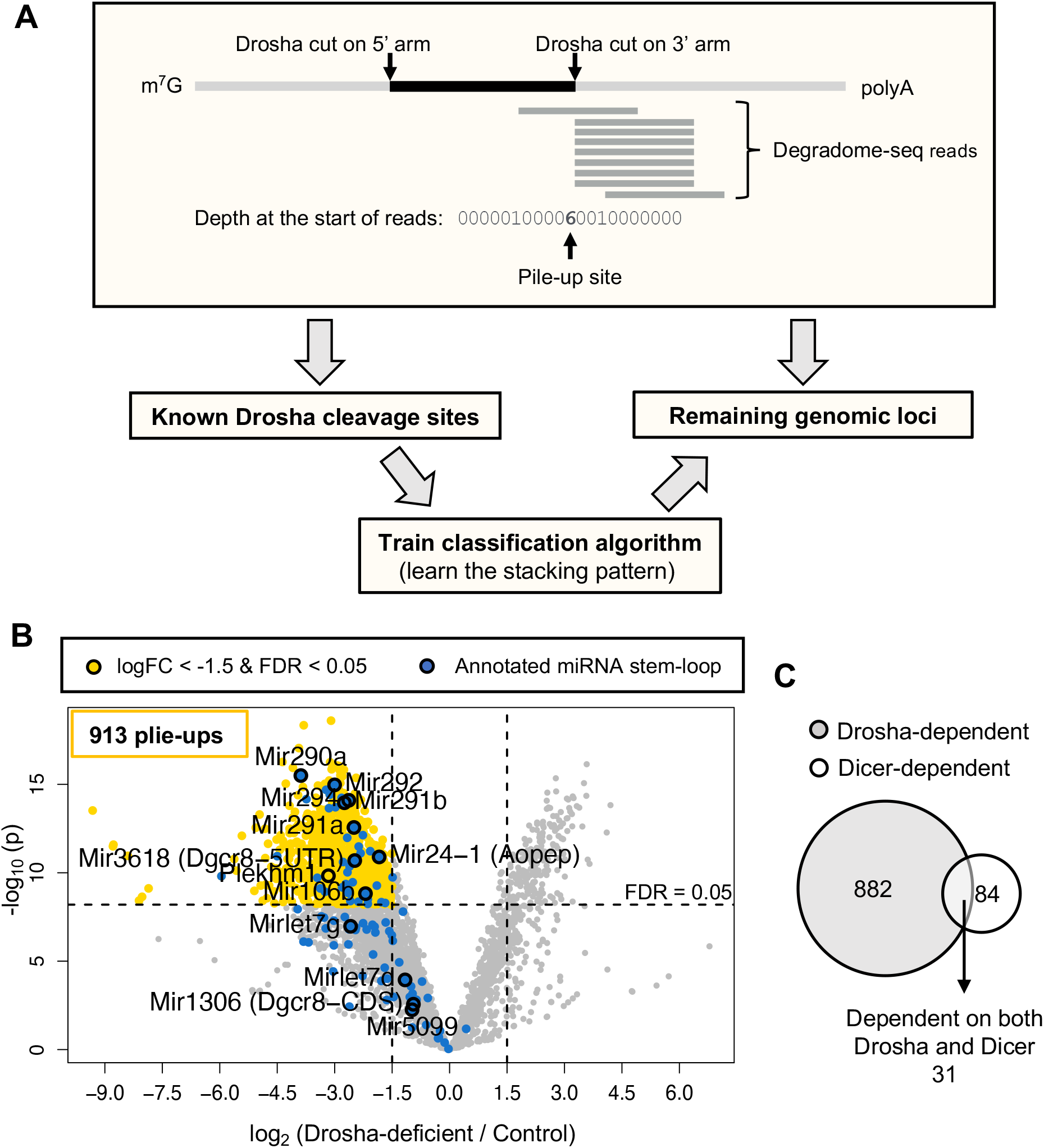
Analysis pipeline for identifying Drosha-dependent cleavage sites captured by Degradome-seq. **(A)** Schematic of the pipeline used to identify pile-up sites in Degradome-seq libraries. For each genomic locus, the depth at the start of the Degradome-seq reads was counted. The pile-up at annotated miRNA stem-loops (i.e. known Drosha cleavage sites) was used to train the classification algorithm for identifying the stacking pattern of Drosha-cleaved RNAs. The trained algorithm was then applied to the remaining genomic loci to identify those with similar stacking patterns. **(B)** Volcano plot of site pile-ups comparing Drosha-deficient and control ESCs. Sites that are significantly decreased (logFC < −1.5 and FDR < 0.05, 913 sites) in Drosha-deficient cells are indicated in yellow. Known cleavage sites on annotated miRNA stem-loops are in are indicated in blue. **(C)** Venn diagram of cleavage sites in ESCs that are dependent on Drosha (decreased in Drosha-deficient cells) versus Dicer (decreased in Dicer-deficient cells).

### Most of the Drosha-dependent cleavage sites are miRNA-independent

A Drosha-dependent pile-up site was identified in the *Plekhm1* mRNA (Figure 2B). Cleavage of *Plekhm1* has previously been shown to be a miR-106b-5p-guided cleavage target and is mediated by the slicer activity of Argonaute2 (22). This also leaves a 5’P on the cleaved RNA (Supplementary Figure S2). Drosha is required for miR-106b biogenesis, and thus loss of Drosha affects *Plekhm1* cleavage indirectly.

To determine if other identified pile-up sites are also miRNA-dependent, we performed Degradome-seq on Dicer deficient ESCs, in which the biogenesis of miRNAs will also be disrupted. Loss of miRNA expression was confirmed following *Dicer1* gene inactivation (Supplementary Figure S3A and B). We generated two independent replicate Degradome-seq libraries each from Dicer-deficient and control ESCs (Supplementary Table S1). Following processing, we identified 4265 pile-up sites in the control cells, of which 115 were lost in Dicer-deficient cells (Supplementary Figure S3C; Supplementary Table S3). Of these, only 31 sites were affected by both Drosha and Dicer deficiency, and presumably via loss of miRNAs (Figure 2C). This suggests that most Drosha-dependent RNA cleavage sites in ESCs is independent of both Dicer and miRNA.

### Inferring the secondary structure of RNAs that are cleaved by Drosha

Drosha usually cleaves stem-loops at two opposing strands to leave an RNase III enzyme signature 3’ 2 nt overhang (37). To understand the nature of Drosha substrates, we first needed to identify the paired cleavage sites, i.e., the precise position of the cleavage on both the 5’ and 3’ arms of a putative stem loop structure. Degradome-seq reveals the cleavage site on the 3’ arm, but not on the 5’ arm. To identify the 5’ cleavage site, we employed sRNA-seq libraries, which capture miRNAs and other by-products of Drosha cleavage, such as miRNA-offset miRNAs (moRNAs) and other sRNAs. It has previously been shown that moRNAs are derived from the sequence immediately adjacent to the mature miRNA, with one end corresponding to the Drosha cleavage site and the other is thought to be processed by non-specific exonucleases (38) (Figure 3A). We reasoned that Drosha cleavage of non-miRNA stem-loops would also result in similar by-products that are captured by sRNA-seq.

**Figure 3.**
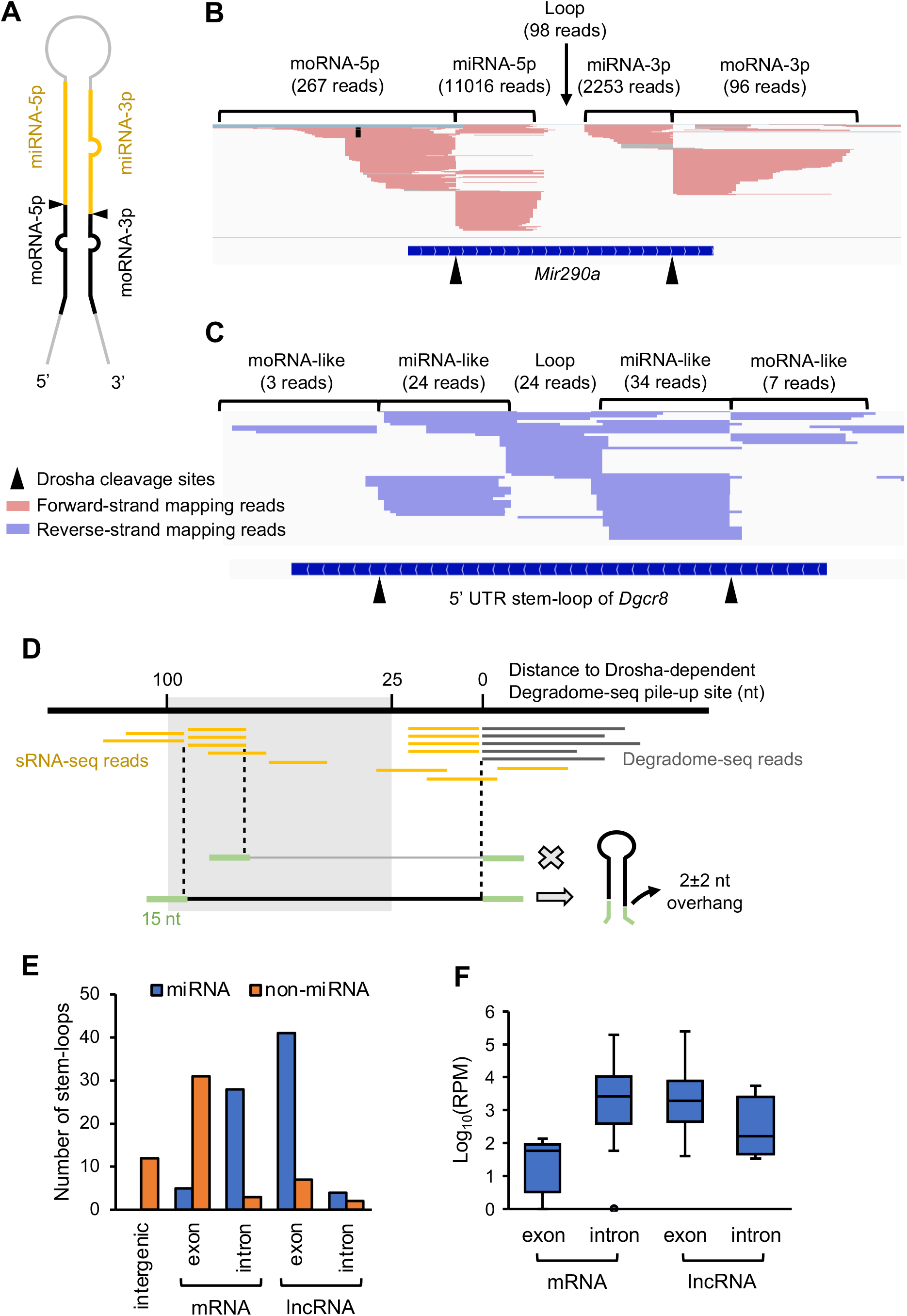
Identifying the boundaries of Drosha-cleaved stem-loops using miRNA and moRNA fragments from sRNA-seq. **(A)** Schematic representation of a miRNA stem-loop structure within the pri-miRNA transcript. Indicated are the position from where the miRNAs (yellow) and moRNA (black) fragments are derived. The Drosha cleavage sites are indicated by the black triangles. **(B)** sRNA-seq reads mapping to the primary *Mir290a* locus. Individual reads are shown. **(C)** sRNA-seq reads mapping to the 5’ UTR stem-loop of *Dgcr8*. Individual reads are shown. **(D)** Strategy for identifying Drosha-cleaved stem-loops from sRNA-seq reads. Reads starting or ending between 25 and 100 nt from the Drosha-dependent Degradome-seq pile-up were collected. If the sRNA-seq reads stack at one of the termini and the depth of that terminus is greater than one RPM, it is considered a potential Drosha cleavage site for the opposite arm of the stem-loop. The RNA sequence between the terminus and the corresponding Degradome-seq pile-up site, flanked by a 15 nt sequence, was then used to predict the stem-loop structure. A stem-loop was considered a Drosha cleavage substrate if the terminus of the sRNA-seq reads and the Degradome-seq pile-up site formed a Drosha signature 3’ 2±2 nt overhang. **(E)** Genomic locations of Drosha-cleaved stem-loops identified in ESCs. Annotated miRNA stem-loops are indicated in blue and non-miRNA stem-loops are indicated in orange. (F) The levels of annotated miRNAs derived from mRNAs and lncRNAs. The levels of each miRNA was obtained from miRBase (56).

We retrieved 38 sRNA-seq libraries of ESCs from the NCBI GEO database (Supplementary Table S4). Low quality reads were first removed from each library, then the libraries were collapsed into unique reads and aligned to the mouse reference genome. Collapsing reads increases the representation of moRNAs and other low-frequency reads in the library to allow for easier identification. We confirmed that moRNAs do indeed map to miRNA loci immediately adjacent to the mature miRNA sequences, with the 3’ end of the moRNA-5p and the 5’ end of the moRNA-3p corresponding to the Drosha cleavage sites on the 5’ and 3’ arms of the pri-miRNA stem-loop (Figure 3B). Distal termini of moRNAs were of varying lengths, consistent with exonuclease-mediated degradation. Similar sRNAs were found at adjacent to Drosha-meditate cleavage site in the stem-loop at the *Dgcr8* mRNA and other non-miRNA Drosha cleavage targets (Figure 3C and Supplementary Figure S4). Here were refer to these as ‘miRNA-like’ and ‘moRNA-like’ sRNAs. Such by-product sRNAs appear to be common to all Drosha substrates and not unique to miRNA precursors.

An algorithm was developed to use these moRNA-like sRNA-seq reads to determine position of the cleavage site on 5p arm of Drosha targets, which were then aligned to the Degradome-seq pile-up sites (Figure 3D). For this, we collected the sRNA-seq reads that started or ended between 25 and 100 nt from the Degradome-seq site, which corresponds to the size of known pre-miRNA stem-loops. Of the Drosha-dependent sites identified by our Degradome-seq of ESCs, 801 exhibited this sRNA read pattern. If the sRNA-seq reads stacked at one of the termini and the depth of that terminus was greater than one RPM, it was regarded as a potential site of Drosha cleavage. The secondary structure of the sequence between the sRNA terminus and the corresponding Degradome-seq pile-up site, flanked by a 15 nt segment, was then predicted (Figure 3D). The flanking sequences were included to capture the entire putative stem-loop structure. A sequence that is predicted to fold into a stem-loop, with termini that formed an RNase III signature 3’ 2 nt overhang, was considered a cleavage target of Drosha (Figure 3D). This correctly identified the 5’ and 3’ Drosha-mediated cleavage sites in 73 miRNA stem-loops out of the 74 miRNAs that are highly expressed in ESCs, demonstrating the effectiveness of the algorithm. We were then able to reassemble 40 non-miRNA stem-loops that are cleaved by Drosha in ESCs (Supplementary Table S5).

Long non-coding (lnc)RNAs are a class of RNA that are longer than 200 nt. They usually possess a 5’ m^7^G cap and a 3’ polyA tail, but they do not encode functional proteins (39). Many miRNA stem-loops are mapped to lncRNA genes, and the level of mature miRNAs derived from these transcripts is relatively high (Figure 3E and F). Thus, these lncRNA are actually pri-miRNAs (Figure 3E). One-third of the miRNA stem-loops were located in mRNA introns, which is consistent with previous studies (40). Only four annotated miRNA stem loops were located within an exon of a mRNA and these miRNAs tended to be expressed at low levels (Figure 3F). Given the low level of mature miRNAs, these exon-located stem loops are unlikely to be functional miRNA precursors. In contrast, most of Drosha-targeted non-miRNA stem-loops were located in the exon of mRNAs (Figure 3E).

### Drosha-cleaved non-miRNA stem-loops are less thermodynamically stable than miRNA precursor stem-loops

Drosha cleavage of exonic stem-loops is expected to destabilize the mRNA (18,19,21). Stem-loops are one of the most common RNA structures (41), and yet Drosha cleaves only some stem-loops but not others. We thus sought to understand the nature of the stem-loops that are recognized and cleavage by Drosha. For this, we systematically characterized and compared the features of miRNA and mRNA stem-loop targets of Drosha. If more than one alternative Drosha cleavage sites were identified for a stem loop, the site with the highest number of sRNA-seq reads was selected for analysis.

We found that mon-miRNA stem-loops had a significantly higher minimum free energy (MFE) compared to miRNA stem-loops, indicating that they were less thermodynamically stable (Figure 4A). The stability can be affected by several properties, including RNA length, base pairing availability, and base pairing composition. We found that non-miRNA stem-loops are slightly shorter but had a significantly higher variance in length compared to the miRNA stem-loops (*F*-test, p < 0.0001). In addition, miRNA stem-loops had a higher base pairing frequency and a longer base pairing stacking (Figure 4B). These results suggest that base pairing plays a significant role in the stability of miRNA stem-loops.

**Figure 4.**
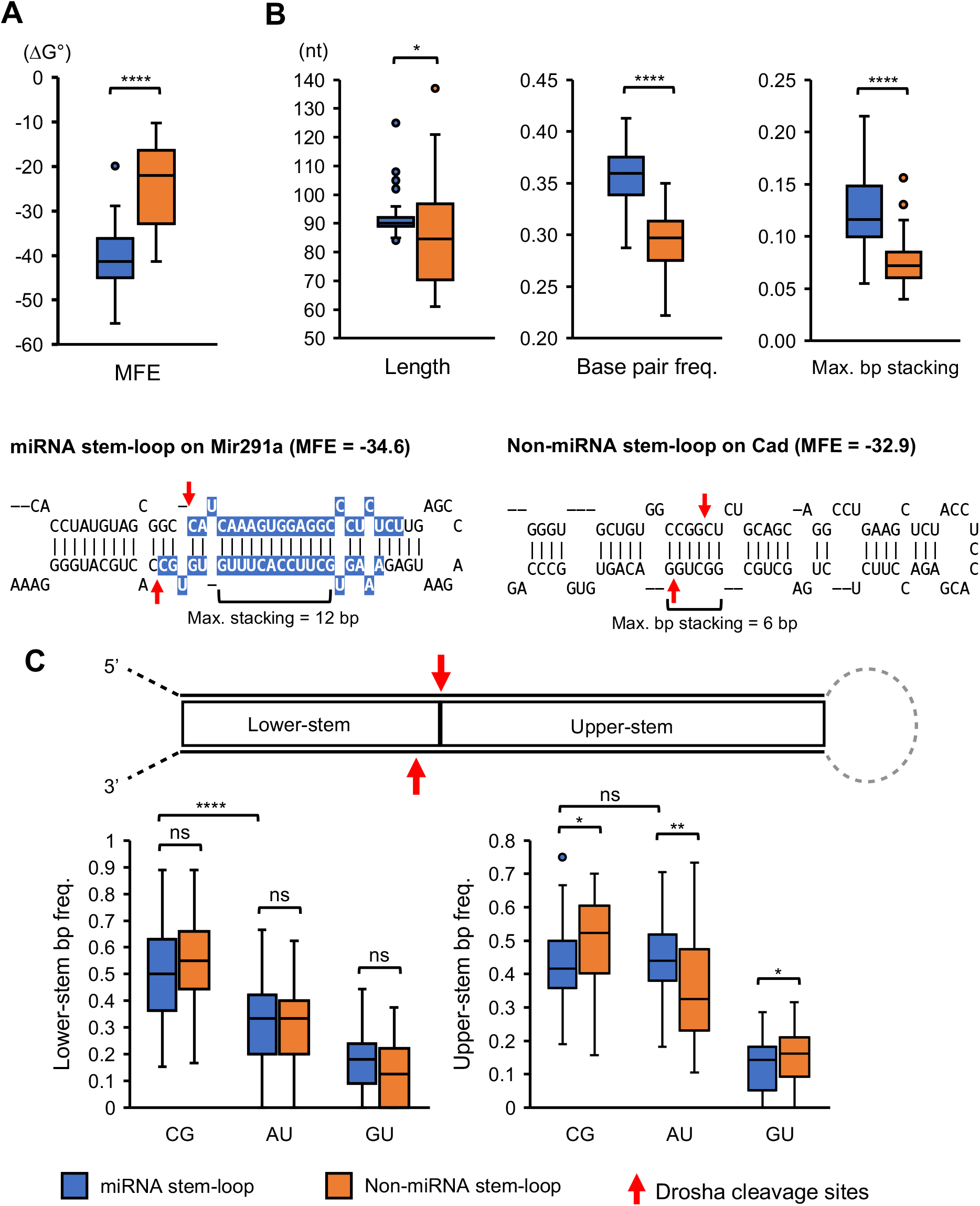
The thermodynamic properties of miRNA and non-miRNA stem-loops. **(A)** Comparison of the MFEs of all Drosha-cleaved annotated miRNA and non-miRNA stem-loops identified in ESCs. **(B)** Comparison of the stem-loop length, base pairing frequency, and maximum stacking of base pairing between Drosha-cleaved annotated miRNA and non-miRNA stem-loops identified in ESCs. The stem-loop in *Mir291a* and *Cad* are shown here to illustrate the typical difference between the miRNA and non-miRNA stem loops. The miRNA sequences are highlighted in blue. The Drosha cleavage sites identified in ESCs are indicated by red arrows. **(C)** Comparison of the base pair composition of lower- and upper-stem of miRNA and non-miRNA stem-loops. The base pair frequency is normalized to the total base pairs in the lower- or upper-stem. The box represents the interquartile range (IQR), the whiskers extend to the most extreme data points within 1.5 times the IQR, and outliers are shown as individual points in boxplots. The median is represented by the line within the box. Statistical testing in **(A), (B)** and **(C)**: * *p* < 0.05; ** *p* < 0.01; *** *p* < 0.0001; **** *p* < 0.00001; ns, not significant (*t*-test).

The composition of base pairing appeared to have less of an impact. Both miRNA and non-miRNA stem-loops exhibited similarly high G-C base pairing and low A-U and G-U base pairing in the lower-stem (Figure 4C). However, only non-miRNA stem-loops exhibited a preference for high G-C base pairing in the upper-stem whereas there was a similar frequency of A-U and G-C base pairing in the upper-stem of miRNA stem loops (Figure 4C). Mature miRNAs originate from the upper-stem. This similar A-U and G-C usage in the upper-stem of miRNA stem loops likely relates to the requirement for sequence diversity in miRNAs. Therefore, the stability of miRNA stem-loops is primarily achieved through extensive base pairing, while non-miRNA stem-loop relies mainly on the G-C base pairing.

### Non-miRNA stem-loops display more alternative structure

An analysis of positional base pairing entropy shows that miRNA stem-loops have low entropy in the stem region, with a rise in entropy in the ssRNA region, including the unstructured flanking region and terminal-loop region (Figure 5A). These suggest that the stem of miRNA stem-loops is unlikely to form alternative base pairing, which likely ensures precise cleavage by Drosha and therefore the sequence of the mature miRNA that is eventually produced. In contrast, the entropy for non-miRNA stem-loops was consistently high, implying the presence of numerous alternative base pairing possibilities (Figure 5A). This is further supported by the high ensemble diversity of non-miRNA stem-loops, which reflects the diversity of secondary structures that a non-miRNA stem-loop can adopt (Figure 5B).

**Figure 5.**
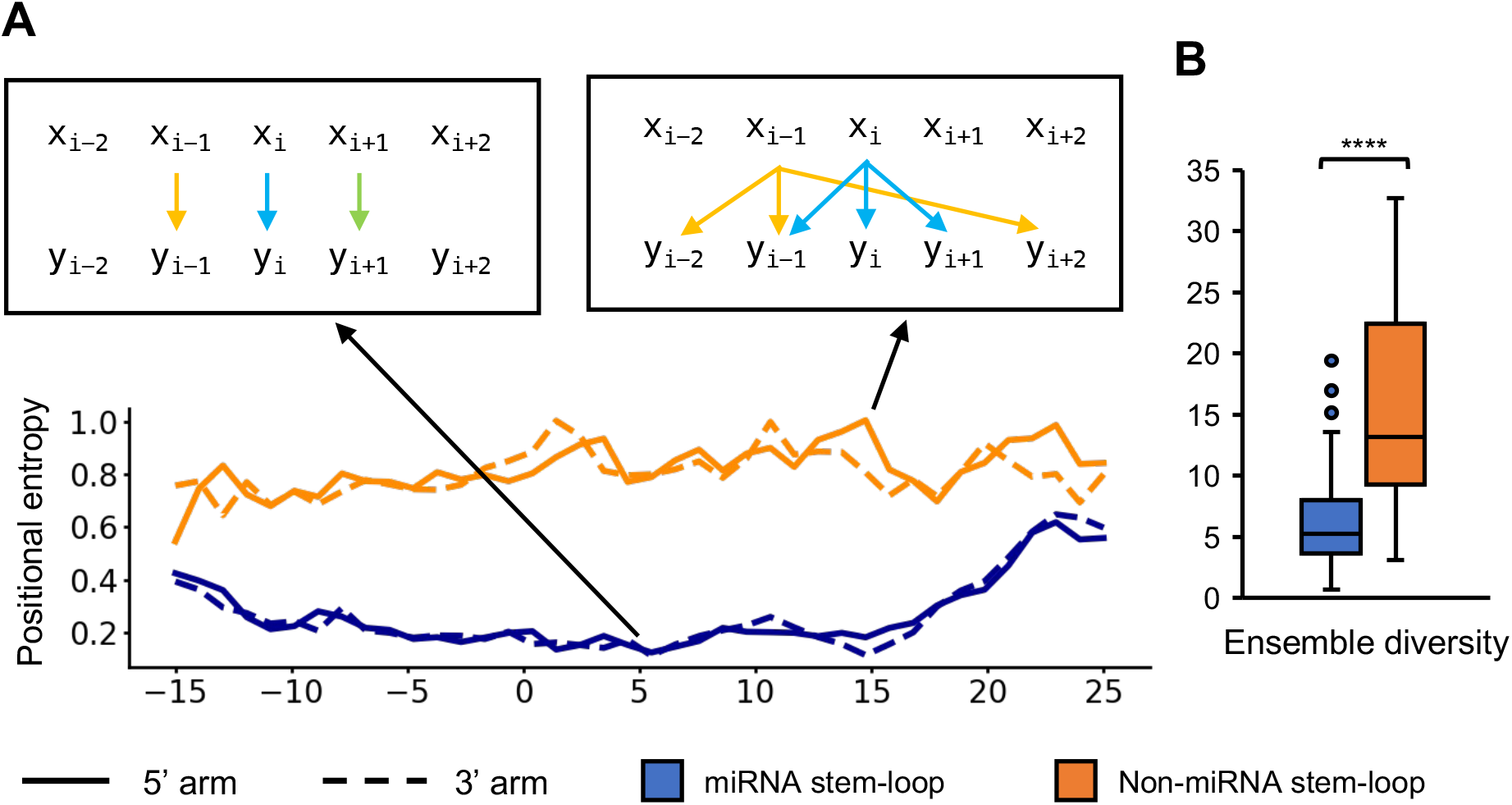
Structure diversity of miRNA and non-miRNA stem-loops. **(A)** Comparing the entropy of Drosha-cleaved annotated miRNA (blue) and non-miRNA (orange) stem-loops identified in ESCs. The 5’ and 3’ arms are plotted in solid and dashed lines, respectively. The x-axis shows the relative position of the stem-loop. Position 0 is the first nucleotide in the upper-stem. **(B)** Comparison of the ensemble diversity of miRNA and non-miRNA stem-loops. The box represents the interquartile range (IQR), the whiskers extend to the most extreme data points within 1.5 times the IQR, and outliers are shown as individual points. The median is represented by the line within the box. Statistical testing: **** *p* < 0.00001 (*t*-test).

### Structural differences between miRNA and non-miRNA stem-loops

We next compared the fine structure of the stem-loops and included 25 nt of the sequence flanking the Drosha-cleaved stem-loop structure. In agreement with the established understanding of the ideal length of miRNA stem-loops (10), the miRNA stem-loops identified were found to be extensively base paired between positions −13 and 22, creating a dsRNA stem of ∼35±1 bp in length (Figure 6A). In contrast, non-miRNA stem-loops were extensively base paired only between positions −13 and 13, resulting in a stem that is on average only ∼26 bp in length (Figure 6B). In addition, miRNA stem-loops have a conserved terminal loop region starting from position 25, while such a region was absent from the non-miRNA stem-loops. This is likely due to the variable stem length of non-miRNA substrates. Instead, a ‘weak’ terminal loop can be observed starting from position 17 (Figure 6B), indicating that some non-miRNA stem-loop may have large terminal loops.

**Figure 6.**
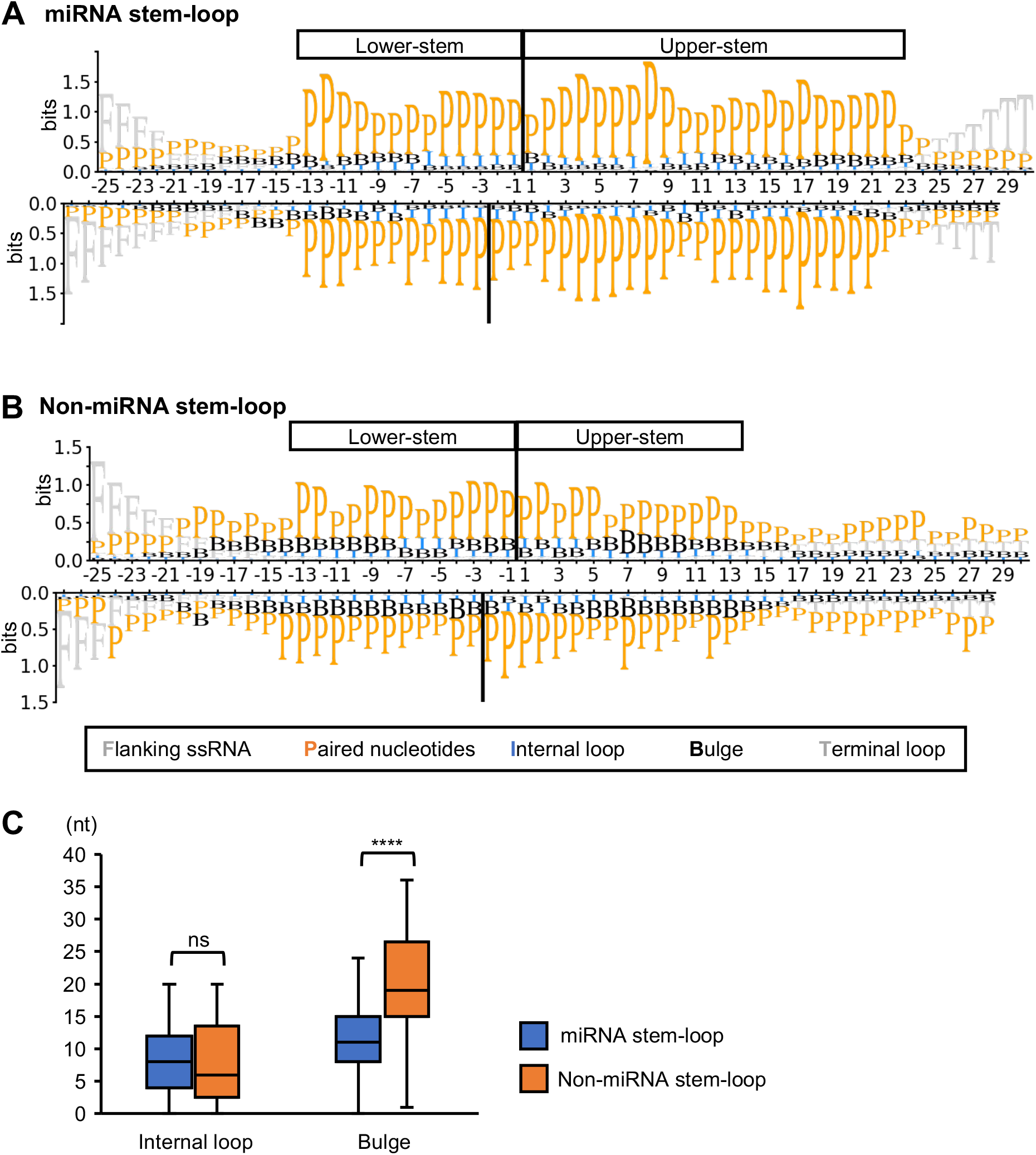
The structural features of miRNA and non-miRNA stem-loops. Information bits plot comparing the structure of Drosha-cleaved **(A)** annotated miRNA and **(B)** non-miRNA stem-loops identified in ESCs. The 5’ arm is shown in the top panels of **(A)** and **(B)**, while the 3’ arm is shown in the bottom panels. The position of the Drosha cleavage site is indicate by solid line. The x-axis indicates the relative position within the stem-loop. P = paired nucleotides; F = flanking ssRNA; I = **i**nternal loop; B = bulge; T = terminal loop. **(C)** Comparison of the number of nucleotides forming internal loops or bulges in miRNA and non-miRNA stem-loops. The box represents the interquartile range (IQR), the whiskers extend to the most extreme data points within 1.5 times the IQR, and outliers are shown as individual points. The median is represented by the line within the box. Statistical testing: **** *p* < 0.00001 (*t*-test).

Non-miRNA stem-loops also displayed more mismatches compared to miRNA stem-loops. Only a few conserved mismatches were observed in miRNA stems, including those at positions −8, −6, 1 and 11 (Figure 6A). The mismatches at positions −6 and 11 have previously shown to improve the accuracy of Drosha processing (10,14). The mismatches at the other two positions may also play a similar role. In contrast, numerous mismatches were observed in non-miRNA stem-loops, most of which formed bulges within the stem (Figure 6B and C).

Despite the numerous structural differences between miRNA and non-miRNA stem-loop, their basal junctions were strikingly similar. A sharp decrease in pairing was observed at position −13 in both types of stem-loops (Figure 6A and B), suggesting that a clear ssRNA-dsRNA junction is a crucial feature for Drosha cleavage. Similar to miRNA stem-loops, Drosha appears to bind to this ssRNA-dsRNA junction in non-miRNA substrates to cleave ∼13 bp away from the junction.

### Non-miRNA stem-loop lacks canonical sequence motifs

Several conserved sequence motifs have been identified in miRNA stem-loop precursors and these have been shown to ensure efficient and accurate binding and processing by Drosha. These motifs include the basal UG motif, the apical UGU motif, the CNNC motif and mGHG motif (10,13). All sequence motifs were found in the Drosha processed miRNA stem-loops identified in ESCs (Figure 7A). The apical UGU motif was found to be the most conserved motif in the miRNAs expressed in ESCs. In contrast, none of these miRNA motifs were detected in non-miRNA stem-loops (Figure 7B).

**Figure 7.**
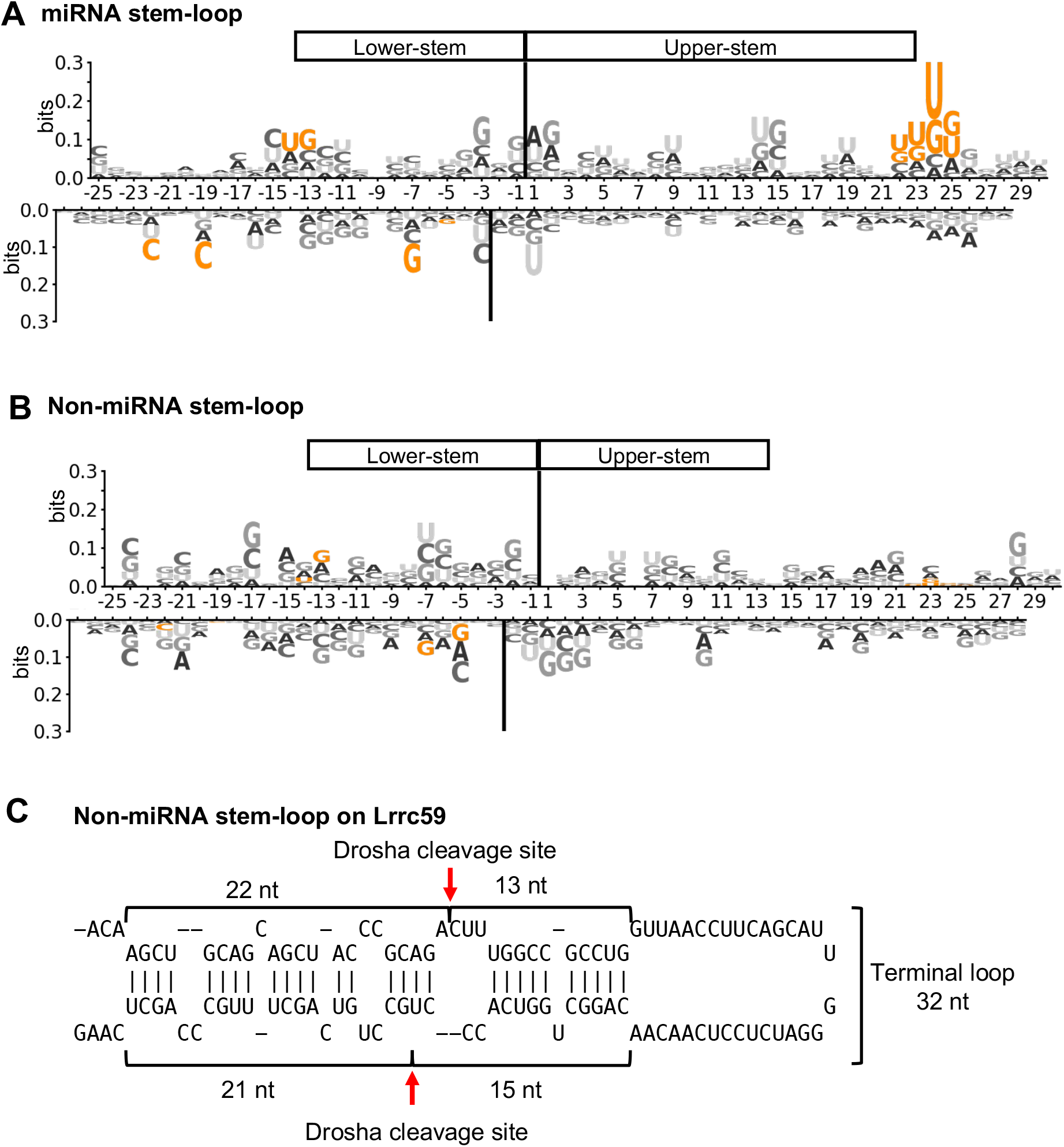
Non-miRNA stem-loops lack known miRNA sequence motifs. Logo of the sequence of Drosha-cleaved **(A)** annotated miRNA and **(B)** non-miRNA stem-loops identified in ESCs. The 5’ arm is shown in the top panels of **(A)** and **(B)**, while the 3’ arm is shown in the bottom panels. The position of the Drosha cleavage site is indicate by solid line. The x-axis indicates the relative position of the stem-loop. Known miRNA motifs are highlighted in orange. **(C)** The non-miRNA stem-loop in *Lrrc59* is likely inversely cleaved due to its large terminal loop and lack of sequence motif. The Drosha cleavage sites are indicted by the red arrows.

Sequence motifs are important for the proper orientation of Drosha when it binds to the stem-loops. Without these motifs, Drosha can bind to the apical junction and cleave the stem-loop inversely (42). While this phenomenon has been demonstrated using pri-miRNA variants *in vitro* with processing assays, whether it actually occurs *in vivo* has been unclear. Our analysis revealed a stem-loop in *Lrrc59* that is inversely cleaved by Drosha in ESCs. This stem-loop is characterized by an unusually large terminal loop of 32 nt. The Drosha cleavage site is ∼13 bp away from the apical junction and has no clear basal junction (Figure 7C), suggesting that Drosha binds to the apical junction and cleaves the stem-loop in an inverse manner. These finding suggest that Drosha recognizes and processes miRNA and non-miRNA stem-loops differently.

## DISCUSSION

Drosha cleaves many non-miRNA stem-loops in mouse ESCs. Our analysis of these non-canonical stem-loop substrates revealed fundamental differences between them and miRNA stem-loops. Specifically, we found that non-miRNA stem-loops are less thermodynamically stable and more likely to fold into alternative structures. Moreover, they lack the sequence and structural motifs normally found in miRNA stem loops that ensure Drosha cleavage at single nucleotide precision. Consequently, Drosha can cleave these non-canonical substrates at positions that are typically inhibited in miRNA stem-loops.

Pri-miRNA stem-loops are typically thermodynamical stable and unlikely to fold into alternative structures. Many non-coding RNAs adopt specific secondary structure to carry out their functions, such as the clove-shaped transfer-RNAs. These RNAs have evolved over time, leading to exceptional thermodynamic stability and low structural diversity, which facilitate their processing and functionality (43,44). Similarly, the stem-loop structure of miRNA plays a crucial role in its biogenesis (3). Thus, like other functional non-coding RNAs, miRNA precursors have a stable structure that enables their efficient and accurate processing by Drosha and Dicer to produce functional miRNAs. However, a stable stem-loop in mRNA would hinder other processes, such as the progression of ribosomes along an mRNA thereby impairing protein expression (45). Therefore, a stable stem-loop would be undesirable in mRNAs, which could explain why non-miRNA stem-loops are generally less stable.

The structure and sequence motifs of miRNA stem-loops are important for accurate binding and processing by Drosha. Drosha needs to bind to the ssRNA-dsRNA junction at the base and cleave ∼11 bp away to produce miRNA intermediates (11,12). However, due to the symmetrical structure of a stem-loop (i.e. ssRNA-dsRNA-ssRNA), Drosha can also bind to the apical junction, which could result in inverse cleavage of the stem, closer to the loop than to the base. To prevent this, several miRNA stem-loop sequences and structure motifs have been found in miRNA stem-loops to encourage binding of Drosha to basal junction and therefore inhibit inverse cleavage. This includes the UGU motif at the apical junction that interacts with Dgcr8, positioning Drosha towards the lower-stem (46), while the CNNC motif interacts with Srsf3 to recruit Drosha to the lower-stem (13,47). The basal UG and mGHG motif also serve as Drosha-interacting motifs, enabling precise binding to the stem-loop (10,12). Recently, a MidMW10 motif, located 10–12 nt away from the Drosha cleavage site in the upper-stem has also been shown to be as essential for preventing inverse cleavage (14). None of these structure and sequence motifs were found in non-miRNA stem-loops that are cleaved in ESCs. Consequently, inverse cleavage of non-miRNA substrates by Drosha, such as the stem-loop in *Lrrc59* that we described, is possible.

A difference between miRNA and non-miRNA stem-loops was anticipated because Drosha cleavage of mRNA stem-loops generally does not produce meaningful quantities of mature miRNAs. Therefore, most features that ensure accurate miRNA production are not necessary for mRNA stem-loop cleavage. However, because these non-canonical stem-loops lack the elements known to enhance Drosha processing, it is likely that they cannot be efficiently cleaved by Drosha alone. It is possible that *trans-* or *cis-*acting elements may facilitate Drosha-mediated cleavage of non-canonical stem-loop substrates to achieve spatiotemporally regulation of specific targets.

It has been shown that post-transcriptional modification of target RNA can affect Drosha processing. N^6^-methyladenosine (m^6^A) on the RNA upstream of the stem-loop has been shown to recruit Drosha to the vicinity of pri-miRNA stem-loops and enhance Drosha processing efficiency (48). Such modification may also be present upstream of non-miRNA stem-loops to ensure Drosha recruitment and to enhance cleavage efficiency. Additionally, the stability of stem-loops may be affected by ADAR enzyme-dependent A-I editing, thereby altering Drosha’s processing efficiency on the stem-loop (49).

Accessory proteins of the microprocessor may also affect non-canonical stem-loop processing. Although the minimal microprocessor complex of Drosha and Dgcr8 alone is sufficient to process pri-miRNA stem-loops, numerous accessory proteins that interact with the complex have been identified. These include the DEAD-box helicases Ddx5)and tumor suppressor p53 that are required for Drosha-mediated processing of a subset of miRNAs (50). Additionally, hnRNP TAR DNA-binding protein 43 (Tdp43) interacts with Drosha to increase its stability and promote processing (51). Such accessory proteins may facilitate the processing non-canonical miRNA stem-loops.

Furthermore, post-translational modifications can also regulate the microprocessor. Acetylation of lysine residues within the N-terminal of Drosha has been found to repress ubiquitin-mediated proteasome decay (52). Deacetylation of Dgcr8 by histone deacetylase 1 (Hdac1) has been found to increase the affinity of Dgcr8 for a subset of pri-miRNA (53). At least 23 phosphorylated amino acids have been found on Dgcr8. Phosphorylation appear to increase the stability of Dgcr8 without affecting its ability to interact with Drosha (54). Similar post-translational modifications can regulate both protein-protein and protein-RNA interactions, thereby affecting Drosha’s processing efficiency on non-canonical stem-loop targets.

Our analysis could be an underestimate of the number of Drosha targets. We focused on stem-loops with two Drosha cleavage sites that are at most 100 nt apart on the same RNA, that is cleavage of both arms of a stem loop. However, it is possible for Drosha to cleave only one strand of the arm (55), and these stem-loop targets would have been excluded from in our analysis. Additionally, structural studies of Drosha suggest that the ssRNA-dsRNA junction is a crucial feature for successful Drosha cleavage (11,12), a junction that can also be present in longer stem-loops or between a pair of sense:antisense transcripts. Whether Drosha can recognize and cleave such structures remains to be determined.

Our study has provided a comprehensive analysis of stem-loops cleaved by Drosha in ESCs, revealing that the non-canonical stem-loop substrates of Drosha differ significantly from pri-miRNA stem-loops. Drosha cleavage-mediated gene repression has been shown to be critical for safeguarding the pluripotency of the stem cells (19-21). Determining how this Drosha cleavage of non-canonical targets is achieved will therefore be critical for to understand the mechanisms regulating stem cell pluripotency, and the knowledge from this study provide a foundation for such future studies into the regulation of Drosha function.

## DATA AVAILABILITY

The data underlying this article are available in NCBI GEO and can be accessed with accession number GSE228299.

## SUPPLEMENTARY DATA

Supplementary Data are available at NAR online.

