## Supplementary Figures for "Non-canonical RNA substrates of Drosha lack many of the conserved features found in primary microRNA stem-loops"

### Supplementary Figure S1

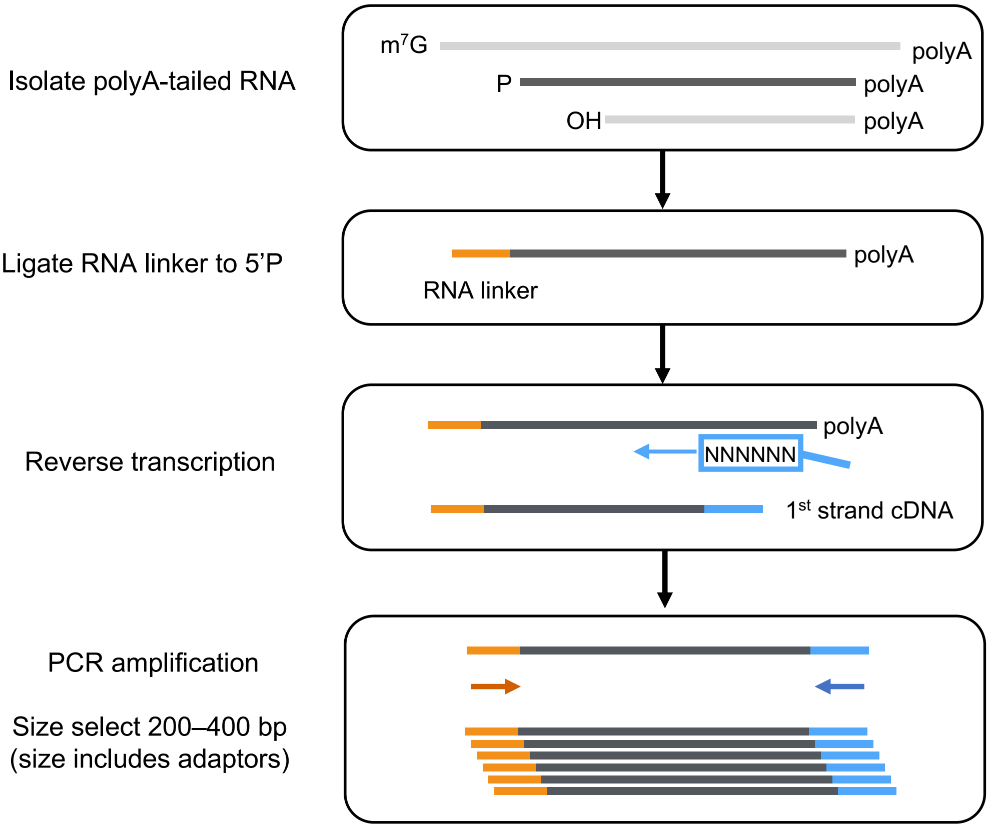


**Supplementary Figure S1.** Degradome-seq workflow. PolyA-tailed RNA was isolated and an RNA linker (in orange) was ligated to fragments containing a 5'phosphate, a hallmark of Drosha cleavage. All RNAs were reverse transcribed with random hexamers linked to a reverse adaptor sequence (in blue). Captured 5'phosphate-containing polyA RNAs were then PCR amplified with primers to the two adaptors, and size selected for high-throughput sequencing on an Ilumina NextSeq 500 platform.

### Supplementary Figure S2

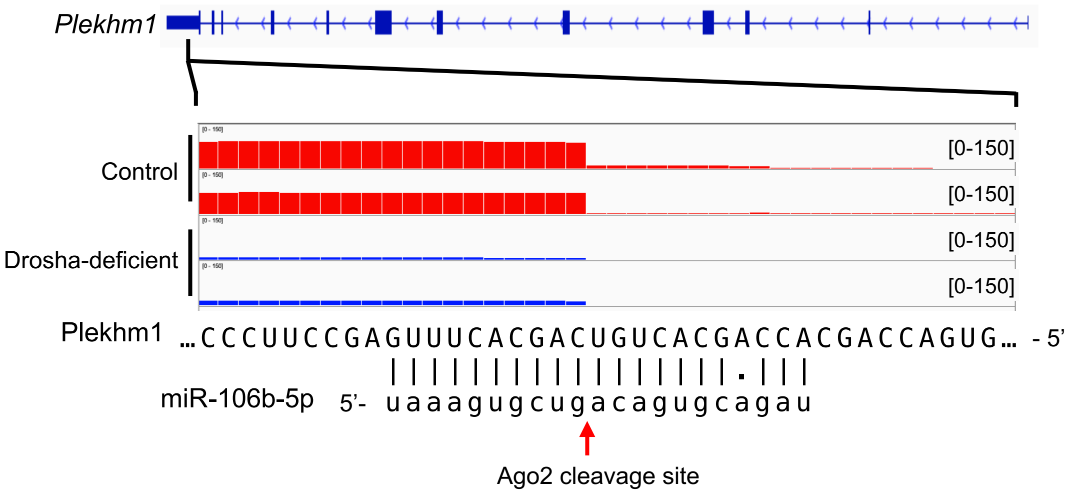


**Supplementary Figure S2.** Ago2-dependent miRNA-guided cleavage of Plekhm1. Shown is the Degradome-seq read pile-up in the *Plekhm1* gene comparing Drosha-deficient (blue) and control (red) ESCs. This is a known Ago2-dependent cleavage site resulting from high complimentary with miR-16-5p sequences. Read depth range is indicated in the square brackets.

### Supplementary Figure S3

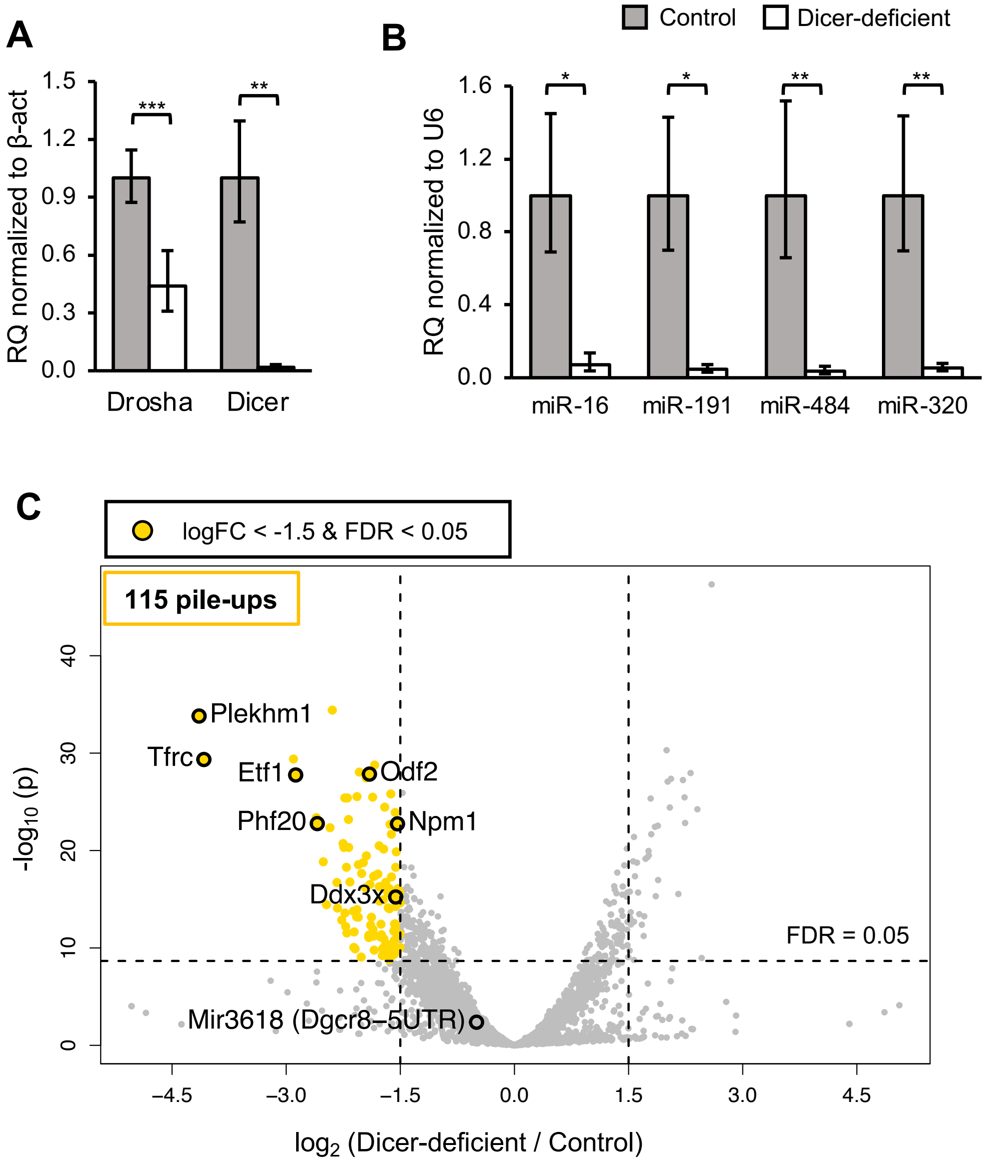


**Supplementary Figure S3.** Identification of Dicer-dependent Degradome-seq sites. **(A)** qRT-PCR analysis of Drosha and Dicer mRNA levels in Dicer-deficient versus control ESCs, normalized to β-actin mRNA. The data shows the mean ± S.E.M of 2 replicates. **(B)** Taqman qRT-PCR analysis for expression of select of mature canonical miRNAs (miR-16 and miR-191) and non-canonical miRNAs (miR-320 and miR-484) in Dicer deficient ESCs, normalized to U6 snRNA. The data shows the mean ± S.E.M of 2 replicates. **(C)** Volcano plot of site pile-ups comparing Dicer deficient and control ESCs. Sites that are significantly decreased (logFC < -1.5 and FDR <0.05, 115 sites) in Dicer deficient cells are indicated in yellow.
Statistical testing in **(B)** and **(C)**: * $p < 0.05$; ** $p < 0.01$ (*t*-test).

### Supplementary Figure S4

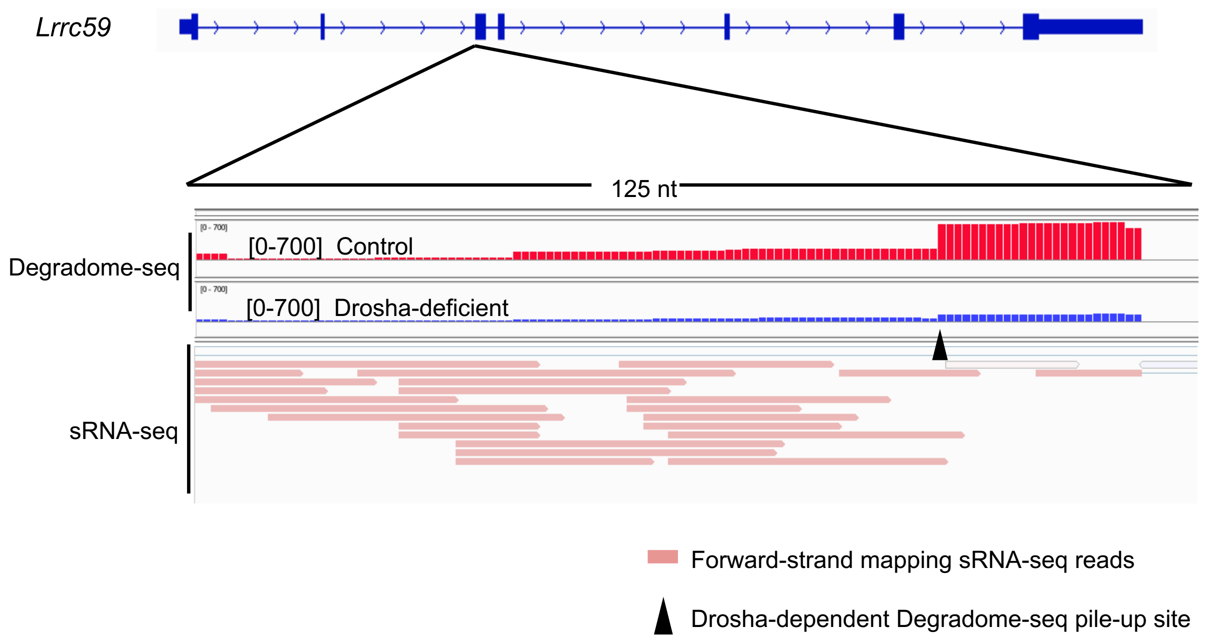


**Supplementary Figure S4.** Alignment of sRNA-seq reads mapping to the Drosha-dependent cleavage site in *Lrrc59* (indicated by black triangle). Depth of Degradome-seq reads comparing Drosha-deficient (blue) and control (red) ESCs is shown in the upper two tracks. The data range for each track is indicated in square brackets. The third track shows a collapsed view of sRNA-seq reads mapping to *Lrrc59*.
